# Heat inactivation of the Severe Acute Respiratory Syndrome Coronavirus 2

**DOI:** 10.1101/2020.05.01.067769

**Authors:** Christophe Batéjat, Quentin Grassin, Jean-Claude Manuguerra, India Leclercq

## Abstract

Supernatants of cells infected with SARS-CoV-2, nasopharyngeal and sera samples containing SARS-CoV-2 were submitted to heat inactivation for various periods of time, ranging from 30 seconds to 60 minutes. Our results showed that SARS-CoV-2 could be inactivated in less than 30 minutes, 15 minutes and 3 minutes at 56°C, 65°C and 95°C respectively. These data could help laboratory workers to improve their protocols with handling of the virus in biosafety conditions.

---

In December 2019, a new coronavirus named SARS-CoV-2 (for Severe Acute Respiratory Syndrome Coronavirus 2) emerged from Wuhan City, Hubei Province in China and quickly became pandemic, spreading in almost all countries in about 2 months. This highly contagious virus has already caused an increasing number of infections and deaths, and the scientific community is facing new challenges to fight the ongoing outbreak. As no specific therapeutics and vaccines are available for disease control, countries lockdown, social distancing and quick detection of cases are currently the main weapons against the virus. Diagnostic and serological tools, by detecting SARS-CoV-2 carriers and immunized recovered patients as soon as possible, are part of this fighting strategy, and necessary to consider a return to normal life. In this context of moving and exponential research, viral inactivation procedures are urgently needed to allow safe experimental laboratory conditions for staff. Amplification of the viral RNA by quantitative RT-PCR is currently the gold standard procedure for diagnosis recommended by the World Health Organization (1). Viral RNA extraction kits sometimes require an initial lysis step at 70°C or more for 5 minutes, but not always. Heat inactivation of the virus is also needed for serum treatment before ELISA and serological assays.

In this study, we tested 3 different inactivation temperatures *i*.*e*. 56°C, 65°C and 95°C, the former being commonly used for inactivation of enveloped viruses (2,3), although higher temperatures can be used for some viruses (4–6). This temperature is also used for elimination of serum complement. We exposed the SARS-CoV-2 to these three different temperatures during various periods of time and tested its infectivity by TCID_50_ method.

A human strain of SARS-CoV-2, isolated from a French patient hospitalized in February 2020, was grown on VeroE6 cells for three passages. The cells were maintained in Dulbecco’s modified Eagle’s medium (DMEM 1X, GIBCO) and supplemented with 5% fetal calf serum (FCS), antibiotics (0.1 units penicillin, 0.1 mg/mL streptomycin, GIBCO) at 37°C in humidified 5% CO_2_ incubator. The SARS-CoV-2 virus was titrated by TCID_50_ method, as described previously (7), except that VeroE6 cells were used and examination for cytopathic effect was performed after 5 days. The virus with a titer of 6.5 log_10_ TCID_50_/mL was adjusted to a final titer of 6 log_10_ TCID_50_/mL in three different kind of media, *i*.*e*. cell culture media DMEM 1X, pooled nasopharyngeal samples of several patients tested negative for SARS-CoV-2, and pooled sera from donors collected before the 1^st^ of January 2020 at a time when no virus circulated in France.

Diluted samples (500 µL) were submitted in triplicate to various temperatures during different times in a calibrated and verified dry water bath, then cooled on ice and tested for infectivity by TCID_50_ method as described above. All experiments were conducted under strict BSL3 conditions.

Mean viral titers obtained for each condition are presented in Tables 1, 2 and 3. Viral inactivation over time are represented in Figure 1.

**Table 1:** Log_10_ TCID_50_ per mL titers obtained after inactivation of infected cell supernatants containing 6 log_10_ TCID_50_/mL of SARS-CoV-2. ND: not detected (below the limit of virus detection which corresponded to 0.67 log_10_ TCID_50_ per ml).

|  | 56°C |  |  |  |  | 65°C |  |  |  |
| --- | --- | --- | --- | --- | --- | --- | --- | --- | --- |
| Time (minute) | 0 | 15 | 30 | 38 | 60 | 0 | 15 | 30 | 60 |
| Cell culture supernatants | 6.6 | 3.23 | ND | ND | ND | 6.6 | ND | ND | ND |

**Table 2:** Log_10_ TCID_50_ per mL titers obtained after inactivation of nasopharyngeal samples spiked with 6 log_10_ TCID_50_/mL of SARS-CoV-2. ND: not detected (below the limit of virus detection which corresponded to 0.67 log_10_ TCID_50_ per ml).

|  | 65°C |  |  |  | 95°C |  |  |  |
| --- | --- | --- | --- | --- | --- | --- | --- | --- |
| Time (minute) | 0 | 5 | 10 | 15 | 0 | 0.5 | 3 | 6 |
| Nasopharyngeal samples | 6.57 | 4.83 | ND | ND | 6.57 | 6.23 | ND | ND |

**Table 3:** Log_10_ TCID_50_ per mL titers obtained after inactivation of human sera spiked with 6 log_10_ TCID_50_/mL of SARS-CoV-2. ND: not detected (below the limit of virus detection which corresponded to 0.67 log_10_ TCID_50_ per ml).

|  | 56°C |  |  |  |  |  |
| --- | --- | --- | --- | --- | --- | --- |
| Time (minute) | 0 | 5 | 10 | 15 | 20 | 30 |
| Sera | 6.2 | 4.87 | 2.57 | ND | ND | ND |

**Figure 1:**
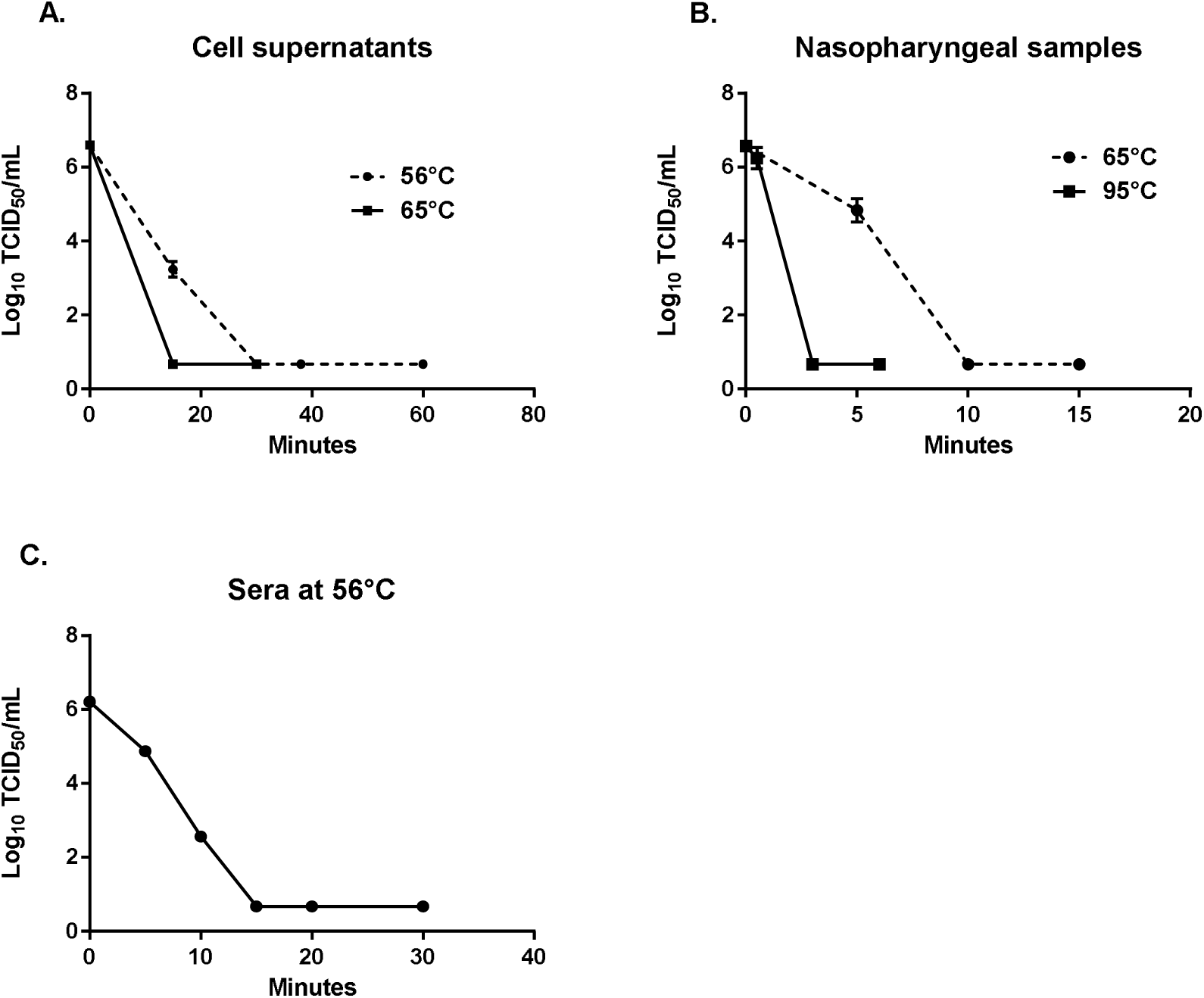
Viral titers in log_10_ TCID_50_/mL obtained after heat inactivation of infected cells supernatants (A), nasopharyngeal samples spiked with SARS-CoV-2 (B) and sera samples from negative donors spiked with SARS-CoV-2 (C). Each condition was performed in triplicate. Each dot represents the mean viral titer and vertical lines represent the standard deviation.

At 56°C, no infectious virus was detected within 30 minutes (Figure 1A and 1C). As expected, increasing the temperature had a negative effect on viral infectivity as no infectious virus was detected within 15 minutes at 65°C (Figure 1A and 1B). At 95°C, 3 minutes were enough to inactivate the virus in nasopharyngeal samples (Figure 1B). The high quantity of infectious virus detected after 30 seconds, *i*.*e*. 6.23 log_10_ TCID_50_/mL, was probably due to the time necessary for the media and the tubes to reach the proper temperature of 95°C inside.

To complement our results, viral RNA from each sample was extracted using a Nucleospin RNA Virus kit (MachereyNagel) for quantitative RT-PCR. RT-PCR targeting two regions of the RdRP gene was carried out using primers and probes developed by the French National Reference Center for Respiratory Infections Viruses with a LightCycler 480 II instrument (8). The E gene was used as a tertiary target for confirmation, following the Charité protocol (9).

The results showed that mean genomic RNA copy number was between 9,9×10^7^ copy genome/5µL and 5,3×10^9^ copy genome/5µL (data not shown). For each condition, the quantity of viral RNA was very stable over time suggesting that viral RNA remained intact in virus particles. At 95°C, the results showed a little decrease of viral RNA quantity over time, resulting in a Cq value raised of about 2.2, potentially preventing SARS-CoV-2 detection for samples with low viral load (10,11).

These results are similar to our previous results obtained with the Middle East Respiratory Syndrome coronavirus (MERS-CoV) (5), which belongs to the same genus of viruses. The SARS-CoV-2 is relatively sensitive to heat inactivation in our laboratory conditions. These data should help laboratory workers to elaborate and improve their protocols for SARS-CoV- 2 experiments, and reinforce our current knowledge on coronavirus survival (12,13).

